# Noninvasive enrichment of circulating tumor biomarkers in a mouse model of diffuse midline glioma using focused ultrasound

**DOI:** 10.64898/2025.12.02.691913

**Authors:** Dingyue Zhang, Yimei Yue, Yan Gong, Leqi Yang, Kevin Xu, Jinyun Yuan, Hong Chen

## Abstract

**Background:** Diagnosing diffuse midline glioma (DMG) through invasive tissue biopsies is challenging due to the tumor’s eloquent location at the pons. Liquid biopsies offer a promising noninvasive alternative; however, their limited detection sensitivity and lack of information on the source of biomarkers pose great challenges. This study aimed to evaluate the feasibility and safety of using focused ultrasound (FUS) with microbubbles to enrich circulating DMG tumor biomarkers in blood and cerebrospinal fluid (CSF) using a mouse model.

**Methods:** Murine DMG cells, transfected with enhanced green fluorescent protein (EGFP) and firefly luciferase (Fluc) genes, were orthotopically injected into the mouse brain. Magnetic resonance imaging was used to guide FUS targeting of the DMG tumor. Droplet digital PCR assays were developed to detect circulating tumor DNA (ctDNA) and RNA (ctRNA) of EGFP and Fluc in blood and CSF samples collected after FUS.

**Results:** FUS enhanced the plasma levels of EGFP ctRNA by 5.4-fold (p=0.0112), compared with liquid biopsy without FUS. CSF EGFP ctDNA was increased by 2.5-fold (p=0.0253), and Fluc ctDNA was increased by 2.6-fold (p=0.0253) with FUS. No brain tissue damage was associated with FUS sonication.

**Conclusions:** This study demonstrated the feasibility and safety of FUS in enriching tumor biomarkers in blood and CSF in a mouse model of DMG. The enrichment ratio for circulating biomarkers depends on the source (plasma vs. CSF), analyte type (ctDNA vs. ctRNA), and individual marker (EGFP vs. Fluc). These findings support the potential future application of FUS to advance the diagnosis of DMG through liquid biopsy.

**Key points:**

1. ddPCR assays were developed to detect plasma and CSF ctDNA and ctRNA in a mouse model of DMG
2. FUS with microbubbles enriched ctDNA and ctRNA in the blood and CSF of a mouse model of DMG
3. The enrichment ratio on circulating biomarkers is dependent on the source (plasma vs. CSF), analyte type (ctDNA vs. ctRNA), and individual marker (EGFP vs. Fluc)

**Importance of the study:** Liquid biopsy via detecting circulating tumor biomarkers holds enormous clinical value in the diagnosis of diffuse midline glioma (DMG), where the eloquent location of the tumor poses significant risks to invasive surgical tissue biopsies. However, the effectiveness of liquid biopsy is hindered by its low sensitivity and the lack of information about the source of biomarkers. This study demonstrated that noninvasive and spatially targeted FUS treatment can release tumor-derived DNA and RNA into the blood and CSF in a mouse model of DMG. This study also found that the enrichment effect of FUS on circulating biomarkers depends on the source (plasma vs. CSF), analyte type (ctDNA vs. ctRNA), and individual marker (EGFP vs. Fluc). This study opens new avenues for advancing noninvasive DMG diagnosis through FUS-enhanced liquid biopsy (sonobiopsy).

## Introduction

Diffuse midline gliomas (DMGs) are indeed a rare type of brain tumor that can arise in various midline structures. Historically, these tumors were primarily referred to as diffuse intrinsic pontine gliomas (DIPGs) because they frequently occurred in the brainstem, specifically the pons, in children^1^. Pons controls vital life functions such as breathing, heart rate, swallowing, and digestion. The eloquent location of tumor in the pons makes surgical tissue biopsies challenging. Repeated surgical biopsies to assess treatment response and recurrence are not feasible due to the eloquent location, the increased risks of infection, bleeding, and other complications^2^. Current standard-of-care disease monitoring approaches rely on radiographic imaging, such as magnetic resonance imaging (MRI)^3^. However, MRI often requires general anesthesia in pediatric patients and cannot provide the molecular characterization of the tumor.

Liquid biopsies analyze specific biomarkers in biofluids like blood and cerebrospinal fluid (CSF), providing a noninvasive/minimally invasive method for diagnosing and monitoring brain tumors. However, using liquid biopsy in DMG patients comes with unique challenges^4^. CSF is a useful source for liquid biopsies due to its proximity to tumor tissue, resulting in higher biomarker concentrations compared to blood^5^. However, obtaining CSF is invasive and often requires general anesthesia in pediatric patients. Though blood collection is less invasive, DMG is a highly diffusive tumor with an often-intact blood-brain barrier (BBB)^6^, which impedes the release of tumor biomarkers into the bloodstream, leading to low biomarker concentration in blood. Additionally, liquid biopsies cannot provide the exact origin of the detected biomarkers, which is essential for monitoring disease progression and improving prognosis.

Transcranial low-intensity focused ultrasound (FUS) in combination with intravenously injected microbubbles has the potential to enrich circulating tumor biomarkers in the blood from the FUS-targeted brain location^7–9.FUS^ combined with microbubbles has been established as a safe and effective technique for noninvasive, spatially targeted, and reversible opening of the BBB^10^. Clinical studies have demonstrated the feasibility and safety of using FUS-induced BBB opening for brain drug delivery for the treatment of adult patients with various brain diseases, including glioblastoma^11,12^, Alzheimer’s disease^13,14^, amyotrophic lateral sclerosis^15^, and Parkinson’s disease^16^. Preclinical studies using mouse models of DMG have reported successful BBB opening for the delivery of therapeutic agents to the tumor^17–23^. One currently active feasibility study is examining the use of this technique with oral etoposide in pediatric patients with progressive DMG (NCT05762419). Building on these technological advancements, we recently demonstrated that FUS combined with microbubbles promoted the release of glioblastoma-derived biomarkers into the bloodstream and improved the detection sensitivity of liquid biopsy^9^. This technique, called sonobiopsy, opened the BBB at spatially targeted brain locations, released tumor-derived biomarkers from precisely defined tumor locations into the blood circulation, and enabled timely detection of biomarkers in the blood to minimize clearance. We showed that sonobiopsy enriched circulating glioblastoma-specific tumor DNAs (EGFRvIII and TERT C228T ctDNAs) in mouse and pig models of glioblastoma^9^. Our first-in-human prospective clinical study in patients with high-grade glioma demonstrated that sonobiopsy is feasible and safe^8^. However, the potential application of sonobiopsy to enhance blood-based liquid biopsies in DMG has not been evaluated. Additionally, the potential of sonobiopsy to enrich brain tumor biomarkers in CSF remains unexplored.

This study aimed to evaluate the feasibility and safety of FUS in enriching circulating biomarkers in blood and CSF in a mouse model of DMG. The mouse DMG model was developed by intracranial injection of murine DMG cells transfected with enhanced green fluorescent protein (EGFP) and firefly luciferase (Fluc). However, the DMG-inherent H3K27M mutation was not present in our DMG cell line. Therefore, we used ctDNA and ctRNA from EGFP and Fluc as surrogate tumor-specific DMG markers. We developed droplet digital PCR (ddPCR) assays for the detection of ctDNA and ctRNA of EGFP and Fluc in blood and CSF samples collected after FUS treatment. We showed that sonobiopsy is feasible and safe for enriching tumor biomarkers in blood and CSF in a mouse model of DMG, with enrichment ratios varying by source (plasma vs. CSF), analyte type (ctDNA vs. ctRNA), and individual marker (EGFP vs. Fluc).

## Materials and Methods

All animal procedures were reviewed and approved by the Institutional Animal Care and Use Committee at Washington University in St. Louis, in accordance with the Guide for the Care and Use of Laboratory Animals and the Animal Welfare Act. A total of 38 mice (18 control and 20 sonobiopsy) were included in the study. The overall experimental scheme is shown in **Figure 1**.

**Figure 1.**
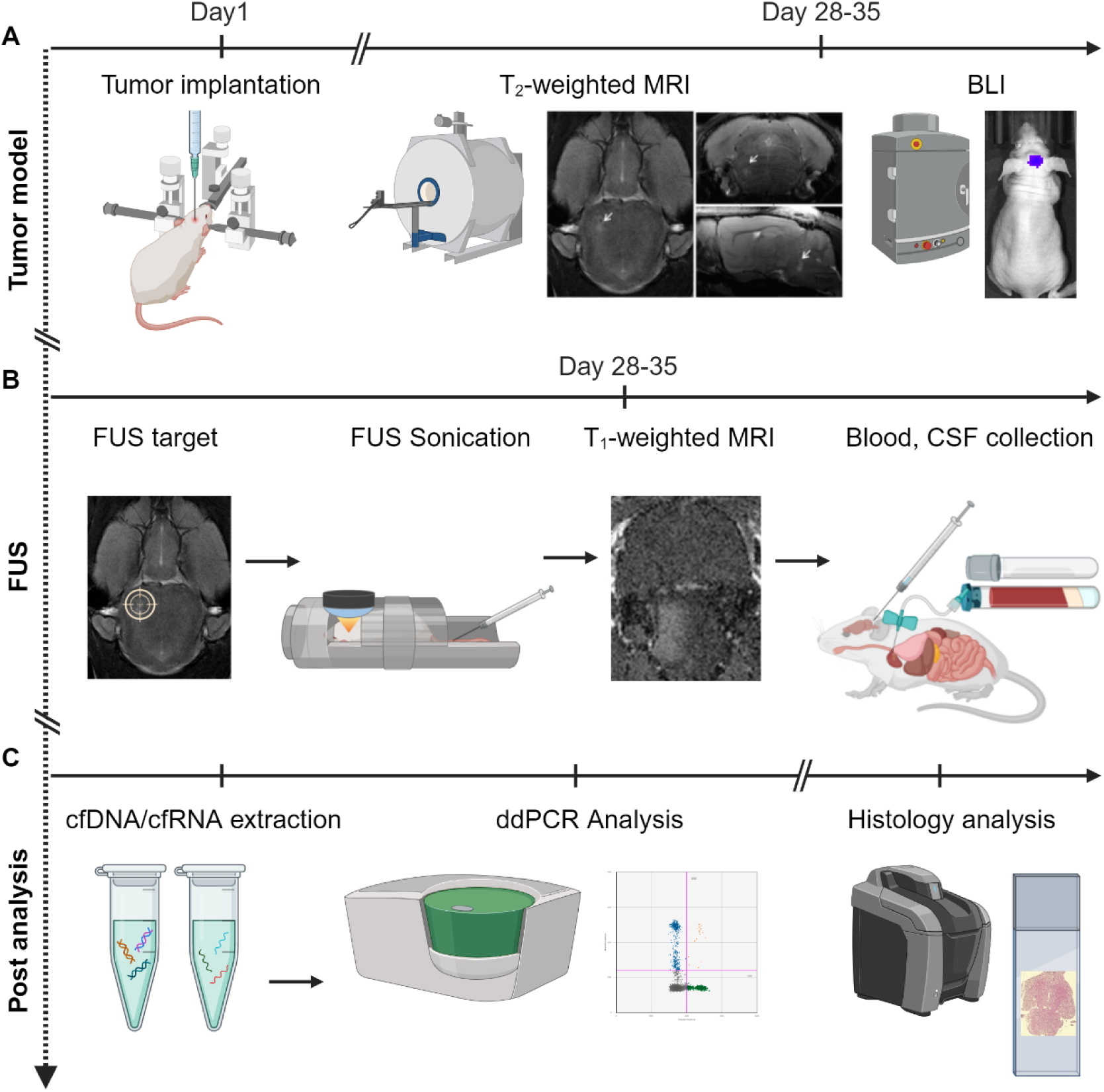
Workflow for sonobiopsy in DMG mice. **A.** Tumor model: Mouse DMG cells were injected into the pontine region of the mouse brain. Tumor growth was confirmed by T_2_-weighted MRI and BLI. **B.** FUS treatment: T2-weighted MRI images were used to align the FUS focal point at the center of the DMG tumor. The FUS transducer was connected to a water balloon for acoustic coupling. Immediately after FUS was turned on, microbubbles and gadolinium contrast were administered via intravenous injection through the tail vein. After FUS sonication, contrast-enhanced T_1_-weighted MRI was obtained to confirm BBB opening. Mice were then sacrificed. Blood, CSF, and brain tissue were acquired. **C.** Post analysis: ddPCR was performed to assess circulating tumor biomarkers in blood and CSF. Histological analysis was used to assess the safety of FUS treatment. Created in BioRender. Chen, H. (2025). https://BioRender.com/swayboj

### DMG tumor cells

Dr. Javad Nazarian from George Washington University kindly provided the murine DMG cells generated from genetically engineered mouse models using RACS/tv-a^24,25^. The murine DMG cells express EGFP and Fluc but not DMG-inherent H3K27M mutation. The cells were cultured at 37°C with 5% CO_2_ in the neurosphere media, which consisted of Neurocult media (Stem Cell Technologies, Vancouver, Canada) supplemented with 10% cell proliferation supplement (Stem Cell Technologies), 2 µg/mL heparin, 20 ng/mL mFGF, 20 ng/mL mEGF, 1% antibiotic-antimycotic, and 50 µg/mL gentamycin.

### DMG mouse model

Immunodeficient mice (strain: NCI Athymic NCr-nu/nu, 6-8 weeks, female, Charles River Laboratory, Wilmington, MA) were used to generate the orthotropic DMG model. Briefly, mice were anesthetized by intraperitoneal injection of a mixture containing ketamine (100 mg/kg body weight) and xylazine (10 mg/kg) in 0.9% saline. A small hole on the skull was created using a microdrill at 1.0 mm lateral to the sagittal suture and 0.8 mm posterior to the lambdoid suture. With a sterile Hamilton syringe, 2×10^5^ cells/2μL in cell culture medium were slowly injected into the pontine tegmentum at 5 mm depth from the inner base of the skull. Mice were monitored daily. Bioluminescence imaging (BLI) was performed by administering luciferin to mice bearing Fluc-expressing tumor cells to monitor tumor growth. Magnetic resonance imaging (MRI) scans using a 9.4T small animal MRI scanner (Bruker, Billerica, MA, USA) were also acquired to confirm tumor growth.

### FUS device

A small-animal MRI-guided FUS system (Image Guided Therapy, Pessac, France) performed FUS sonication following a procedure described in our previous publication^9^. The system consisted of an MRI-compatible FUS transducer (Imasonic, Voray sur l’ognon, France) and a 9.4T small animal MRI scanner (Bruker, Billerica, MA). The FUS transducer was made of a 7-element annular array with a center frequency of 1.5 MHz, a radius of curvature of 20 mm, and an aperture of 25 mm. The FUS transducer’s axial and lateral full widths at half maximums (FWHM) were 5.5 mm and 1.2 mm, respectively. Meanwhile, the acoustic pressure reported in this study was calculated by considering the mouse skull attenuation (18% reduction from the pressure measured in water)^26^. The transducer was connected to a water balloon filled with deionized and degassed water and coupled to the mouse head with degassed ultrasound gel.

### Sonobiopsy procedure

Approximately 4-5 weeks after intracranial tumor implantation, mice were randomly assigned to control (collect blood and CSF without FUS, FUS-group) or sonobiopsy (collect blood and CSF after FUS, FUS+ group). To ensure proper control of the study, both the FUS+ and FUS-groups were exposed to 1.5% isoflurane for around 1 hour and the effect of isoflurane on BBB permeability and tumor biomarker release would be minimal and consistent across both groups^27,28^. The sonobiopsy procedure involved three main steps (**Figure 1B**): (1) FUS target planning; (2) FUS sonication; (3) blood, CSF, and brain tissue collection following procedures like our previous report with some modifications^9^. Briefly, the FUS target planning was performed under the guidance of T_2_-weighted MRI images to align the FUS focal point at the center of the DMG tumor. Before FUS treatment, contrast-enhanced T_1_-weighted MRI scans were performed after intravenous administration of the MRI contrast agent gadoterate meglumine (dose of 1 mL/kg, Gd-DOTA; Dotarem, Guerbet, Aulnay sous Bois, France) to assess the BBB permeability of the DMG tumor before FUS sonication. After the MRI scan, FUS sonication was turned on, followed by intravenously administering microbubbles (Definity, Lantheus Medical Imaging, North Billerica, MA, USA) at a dose of 100 μL/kg. FUS sonication used the following parameters: frequency: 1.5 MHz, derated pressure in tissue: 0.55 MPa, pulse repetition frequency: 5 Hz, duty cycle: 3.35%, pulse length: 6.7 ms, and treatment duration: 3 minutes. FUS sonication targeted nine (3×3) evenly spaced points within a square pattern, with 1 mm spacing between two adjacent points, ensuring tumor volume coverage. Following sonication, contrast-enhanced MRI scans were performed again to evaluate the BBB permeability changes due to FUS sonication. About 10 min after MRI scans (∼20 min after FUS), whole blood (∼500 µL) was collected via cardiac puncture, and CSF was collected via cisterna magna. For FUS-group mice, the samples were collected after exposure to isoflurane for approximately 1 hour. The collected blood was stored in BD Vacutainer K_2_ EDTA tubes. Within 4 hours of collection, blood and CSF tubes were centrifuged at 3000×*g* for 10 minutes at 4 °C to separate the plasma from the hematocrit and remove cell debris from CSF. Plasma and CSF aliquots were put on dry ice immediately for snap freezing and stored at -80°C subsequently for later downstream analysis.

### Quantification of BBB opening with contrast-enhanced MRI

The BBB permeability changes by FUS were quantified based on contrast-enhanced MR images using our previously reported method^9^. Briefly, two circular regions of interest (ROIs) were drawn on the MRI on the FUS-treated site and the contralateral untreated site. Subsequently, voxels within the FUS-treated ROI were identified as indicative of BBB opening if their intensity exceeded the mean intensity of the untreated ROI by 3 standard deviations. Then, the total voxels representing BBB opening were computed by summing the identified voxels across all brainstem sections along the path of the FUS beam.

### Plasma and CSF biomarker analysis using ddPCR

Custom sequence-specific primers and fluorescent probes were designed and synthesized for EGFP and Fluc detection. The forward and reverse primer sequences for EGFP are 5’-GACCACTACCAGCAGAACACC -3’ and 5’-CCAGCAGGACCATGTGATCG -3’, respectively. The EGFP probe sequence is 5’-CCGACAACCACTACCTGAGCACCCAGTC -3’ with the hexachlorofluorescein (HEX) fluorophore and the Black Hole Quencher 1 (BHQ1). The forward and reverse primer sequences for Fluc are 5’-CCAACATCTTCGACGCAGGTG -3’ and 5’-TGACTGGCGACGTAATCCACG -3’, respectively. The Fluc probe sequence is 5’-CCGACGATGACGCCGGTGAACTTCC -3’ with the 6-carboxyfluorescein (6-FAM) BHQ1.

cfRNA/cfDNA was extracted from mouse plasma and CSF using plasma/serum RNA/DNA purification mini kit (Norgen Biotek, Thorold, Ontario, Canada) per manufacturer’s instructions. A high-capacity cDNA reverse transcription kit (ThermoFisher Scientific, Waltham, MA, USA) was used to generate cDNA from cfRNA. cfDNA and cDNA (cfRNA) were pre-amplified for EGFP and Fluc using a Q5 hot start high-fidelity master mix (New England Biolabs, Ipswich, MA, USA). Preamplified products were directly used for further ddPCR reactions. ddPCR reactions were conducted using Bio-Rad Q200X according to the manufacturer’s instructions (Bio-Rad, Hercules, CA, USA). Data were acquired on the QX600 droplet reader (Bio-Rad) and analyzed using QX Manager Software (Bio-Rad). All results were manually reviewed for false positives and background noise droplets based on negative and positive control samples. EGFP and Fluc ctRNA/ctDNA concentrations (copies/µL plasma or copies/µL CSF) were calculated by multiplying the concentration (provided by QX Manager Software) by elution volume, divided by the input plasma volume used during cfDNA/cfRNA extraction.

### Safety evaluation by histological analysis

After blood collection, the mouse brains were extracted after transcardial perfusion with phosphate-buffered saline. Mouse brains were fixed in 4% paraformaldehyde, dehydrated in 30% sucrose, and embedded in optimum cutting temperature at -20 °C. The brain slices were transversely sectioned into slices with a thickness of 10 μm using a Leica CM1860 cryostat (Leica Biosystem Inc., Buffalo Grove, IL). Slices were stained with hematoxylin and eosin (H&E) to assess any damage to the brain tissue. The stained brain slices were imaged using a microscope (Keyence, Osaka, Japan). After color deconvolution, the microhemorrhage was detected based on the pixel hue threshold using the QPathv0.5.0. The microhemorrhage density was calculated as the percentage of positive-pixel area over the total tumor area.

### Statistical analysis

Statistical analysis was performed in Graphpad (Prism) (Graphpad, Boston, MA, USA). Paired parametric t-tests compared the pre-FUS and post-FUS T_1_-weighted contrast-enhanced volume. The Mann-Whitney test was used to compare the level of tumor-specific biomarkers in the control and FUS treatment groups. Unpaired parametric *t*-tests were used for histology data analysis. *P* values are two-tailed unless otherwise specified.

## Results

### The orthotopic mouse model recapitulated DMG

T_2_-weighted MRI images confirmed the growth of DMG tumors, which was indicated by the presence of hyperintense pixels on the image (**Figure 2A**). Contrast-enhanced T_1_-weighted MRI scans did not show significant contrast enhancement, indicating poor BBB permeability (**Figure 2B**). BLI verified the growth of tumor cells in the mouse brain (**Figure 2C**). Ex vivo H&E staining displayed a diffuse tumor (**Figure 2D**). Collectively, these data indicate that this mouse model recapitulated the intact BBB and diffusive tumor observed in clinical DMG patients.

**Figure 2.**
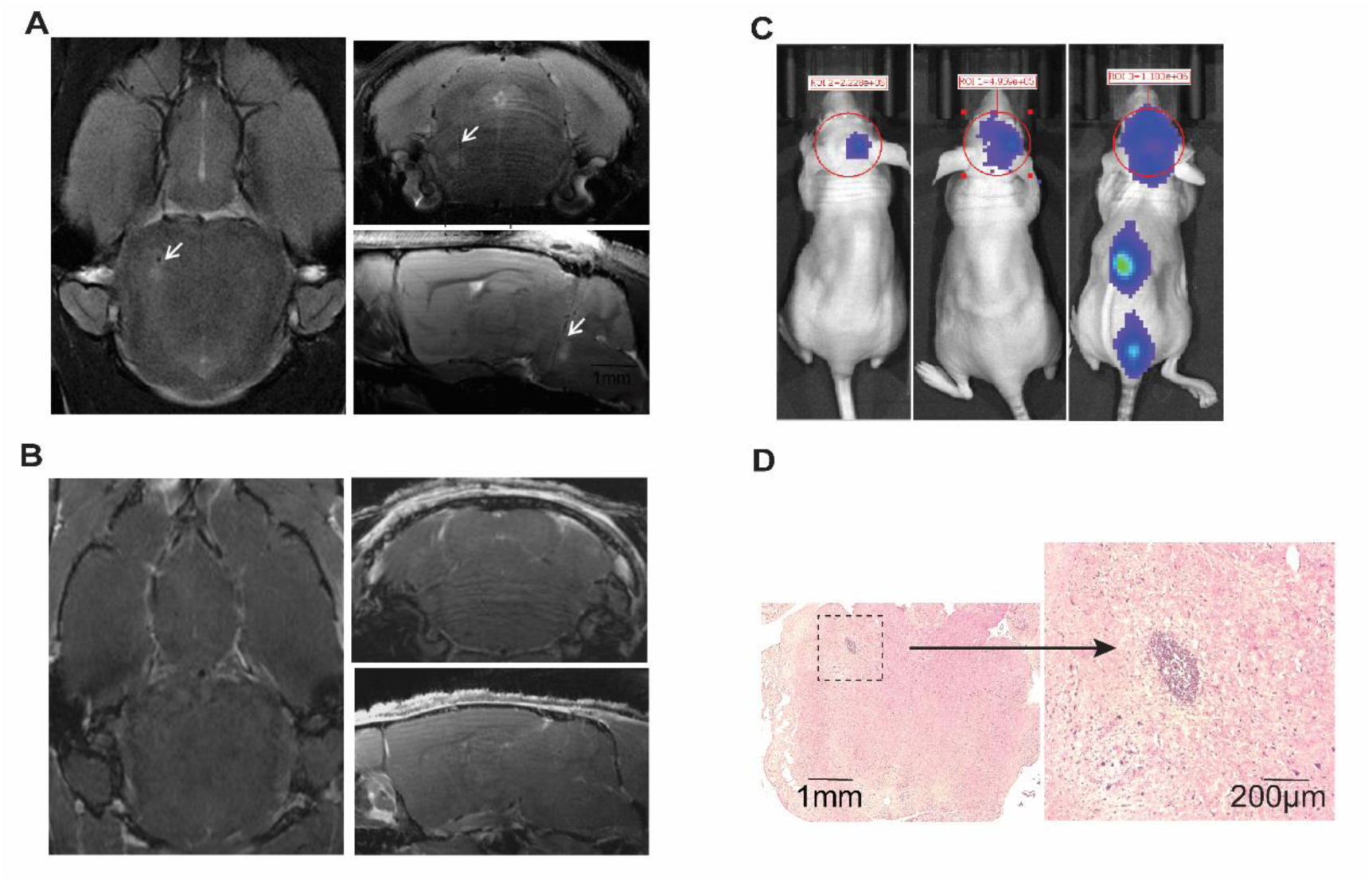
Characterization of DMG mouse model. A. Coronal, axial, and sagittal scan of T_2_-weighted MRI. The tumor is pointed out with a white arrow. B. Coronal, axial, and sagittal scan of contrast-enhanced T_1_-weighted MRI. The tumor does not show enhancement. C. BLI signal of three representative DMG mice. d. Histological analysis of DMG tumor in the mouse poutine area with H&E staining.

### FUS combined with microbubble opened the BBB

Contrast-enhanced T_1_-weighted MRI acquired right before and after FUS sonication shows enhanced pixel intensity at the FUS-targeted brain region post-FUS, indicating increased permeability of the BBB (**Figure 3A**). Quantification of the pixels with enhanced intensity confirmed that FUS significantly enhanced the permeability of the DMG tumors by 102-fold from 0.10 ± 0.02 mm^3^ to 10.45 ± 1.32 mm^3^ (n=18, p<0.0001, paired *t*-test; two mice were excluded from this analysis due to technical issues with MRI; **Figure 3B**).

**Figure 3.**
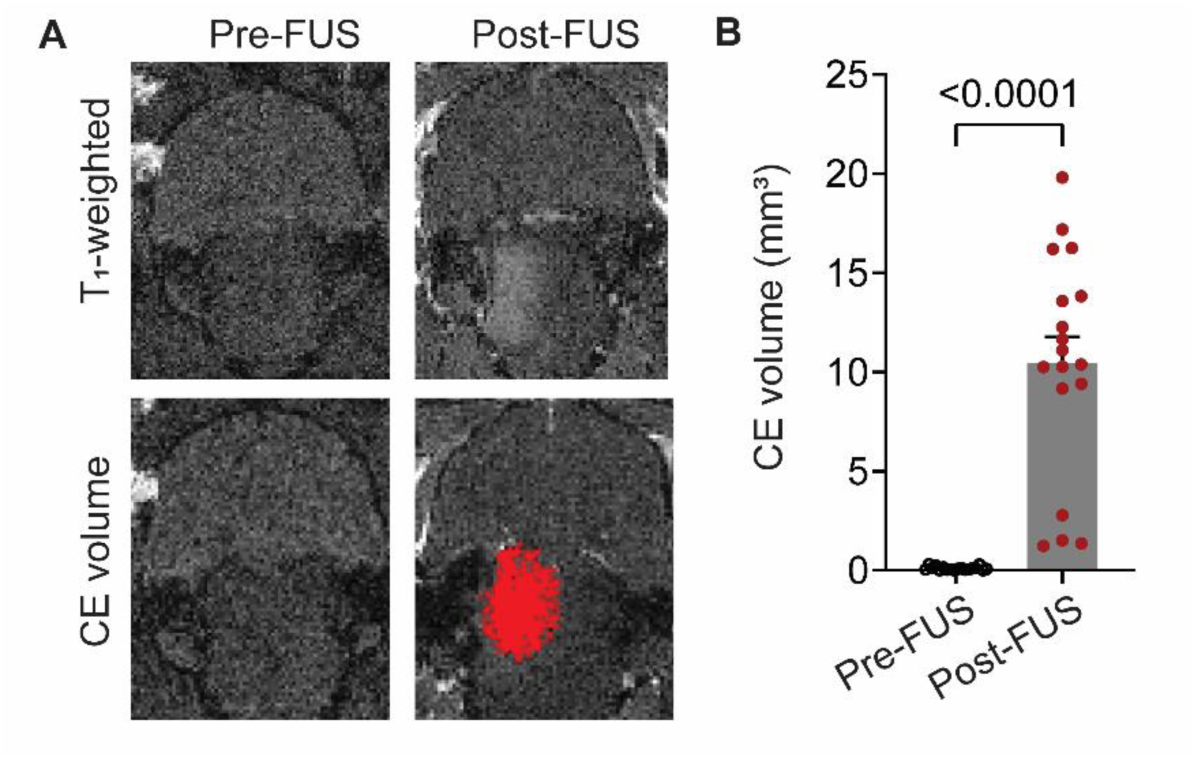
Successful FUS-induced BBB opening in a mouse model of DMG. **A.** Representative contrast-enhanced T_1_-weighted MRI before and after FUS sonication. **B.** FUS significantly increased the contrast-enhanced (CE) volume at the target brain tumor location. The bar graph represents the mean ± standard error of the mean (SEM). Paired *t*-test, comparing post-FUS with post-FUS.

### ddPCR assays were developed for tumor biomarker detection

EGFP and Fluc ddPCR assays were validated with DNA **(Figure 4A)** and cDNA (RNA) **(Figure 4C)** from parent DMG cells. To obtain the limit of blank (LoB) and limit of detection (LoD) of the ddPCR assays, EGFP and Fluc were assayed with serial dilutions of DNA **(Figure 4B)** and cDNA **(Figure 4D)**. LoB was 1.32 copies/rxn (number of copies per reaction tube) for EGFP and 2.08 copies/rxn for Fluc, respectively. LoD was 3.21 copies/rxn for EGFP and 3.22 copies/rxn for Fluc, respectively.

**Figure 4.**
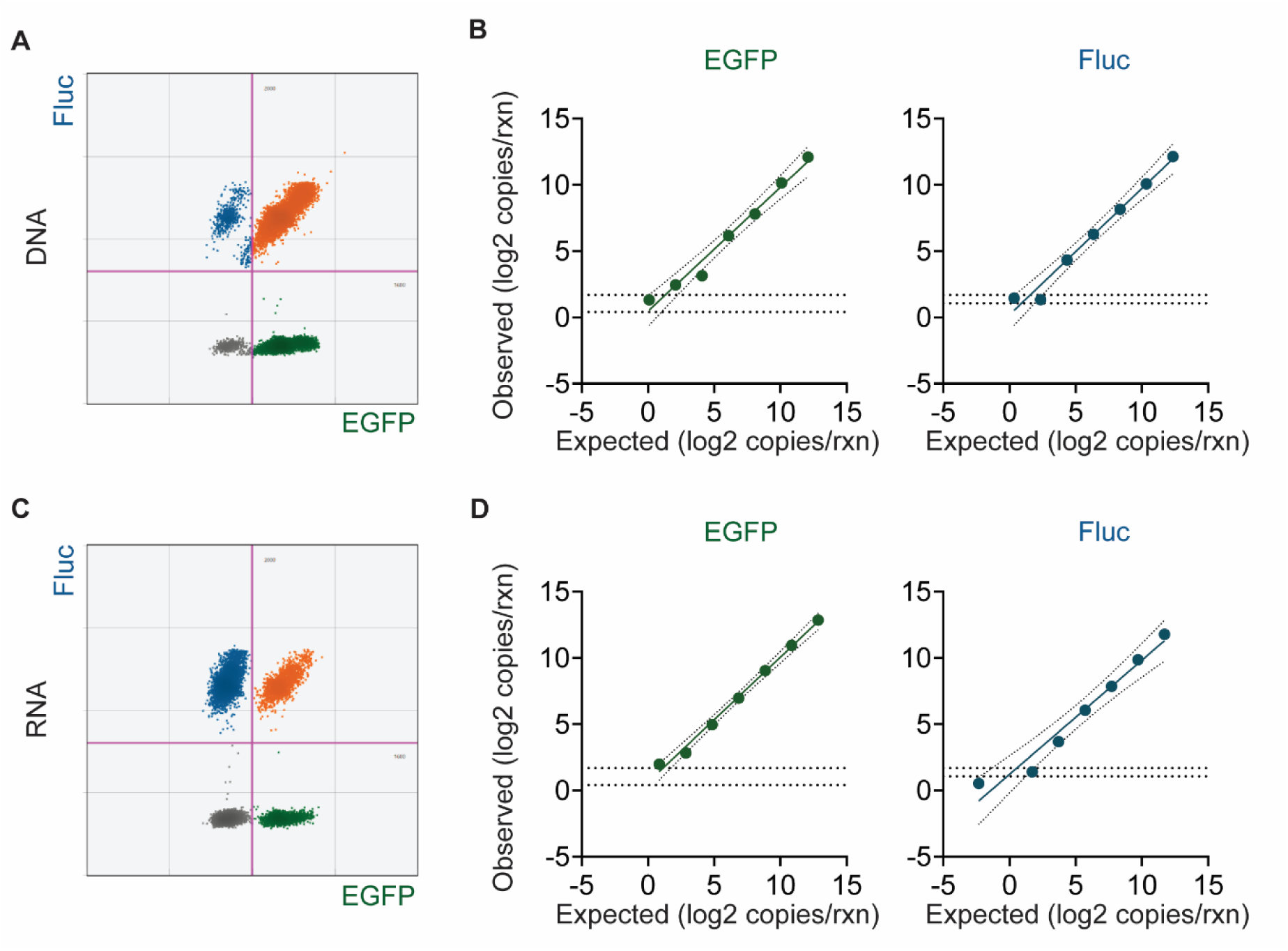
Validation of custom ddPCR primers and probes for detecting DNA and RNA for EGFP and Fluc in DMG cells. **A.** ddPCR detected both EGFP and Fluc in genomic DNA (DNA) from DMG cells. **B.** ddPCR detection of EGFP and Fluc with serial dilutions of DNA **C.** ddPCR detected both EGFP and Fluc in cDNA (RNA) from DMG cells. **D.** ddPCR detection of EGFP and Fluc with serial dilutions of cDNA (RNA). The limit of blank (LoB, lower dashed line) is plotted. LoB = mean of blank + 1.645 × standard deviation of the blank. The limit of detection (LoD, upper dashed line) is also plotted. LoD = LoB + 1.645 × standard deviation of the lowest concentration sample above the blank.

### FUS enriched circulating tumor biomarkers

Blood and CSF samples collected ∼20 minutes after FUS sonication were processed and analyzed by ddPCR. Sonobiopsy (FUS+) notably enhanced the detection of tumor-specific ctDNAs and ctRNAs compared to conventional liquid biopsy (FUS-). FUS+ increased the plasma levels of EGFP ctDNA by 2.0-fold (p=0.806), and EGFP ctRNA by 5.4-fold (*P* = .0112), compared with FUS-**(Figure 5A–D**, **Table 1)**. Plasma Fluc ctRNA was increased by 5.3-fold (*P* = .1542) with FUS treatment **(Figure 5A–D**, **Table 1)**. Furthermore, FUS+ markedly elevated CSF levels of EGFP ctDNA by 2.5-fold (*P* = .0253), EGFP ctRNA by 2.3-fold (*P* = .1942) **(Figure 5E–H**, **Table 1),** compared with FUS-. CSF Fluc ctDNA was increased by 2.6-fold (*P* = .0253), and Fluc ctRNA was increased by 4.6-fold (*P* = .3473) with FUS sonication **(Figure 5E–H**, **Table 1)**.

**Figure 5.**
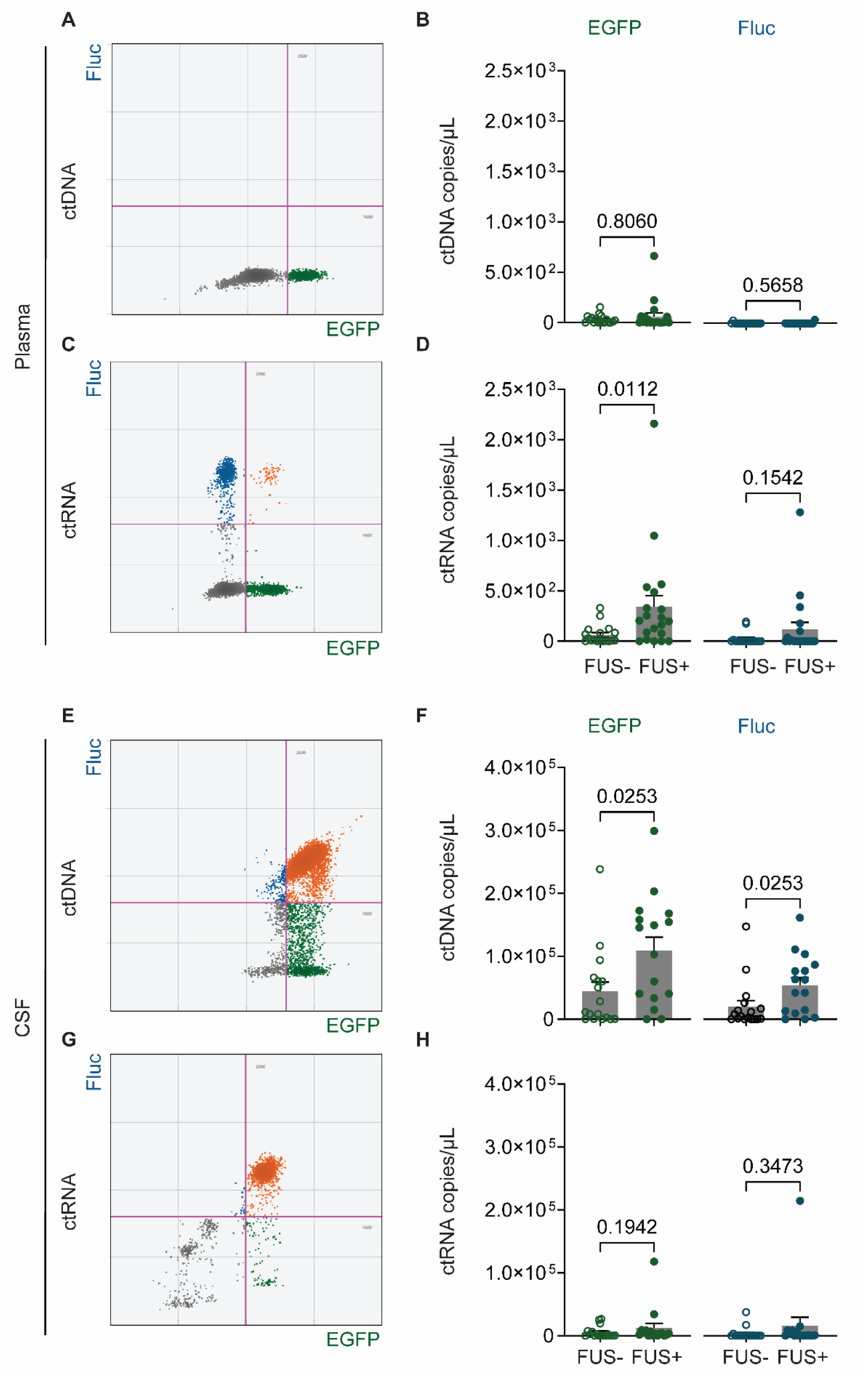
Sonobiopsy enriched blood plasma and CSF biomarkers. **A**. Represent ddPCR plot for EGFP ctDNA and Fluc ctDNA in the plasma. **B**. The normalized amount of EGFP ctDNA and Fluc ctDNA in the plasma. **C**. Represent ddPCR plot for EGFP ctRNA and Fluc ctRNA in the plasma **D**. The normalized amount of EGFP ctRNA and Fluc ctRNA in the plasma. **E**. Represent ddPCR plot for EGFP ctDNA and Fluc ctDNA in the CSF. **F**. The normalized amount of EGFP ctDNA and Fluc ctDNA in the CSF. **G**. Represent ddPCR plot for EGFP ctRNA and Fluc ctRNA in the CSF. **H**. The normalized amount of EGFP ctRNA and Fluc ctRNA in the CSF. The bar graph shows the mean ± standard error of the mean (SEM). Mann-Whitney test, comparing FUS+ with FUS-.

**Table 1.**
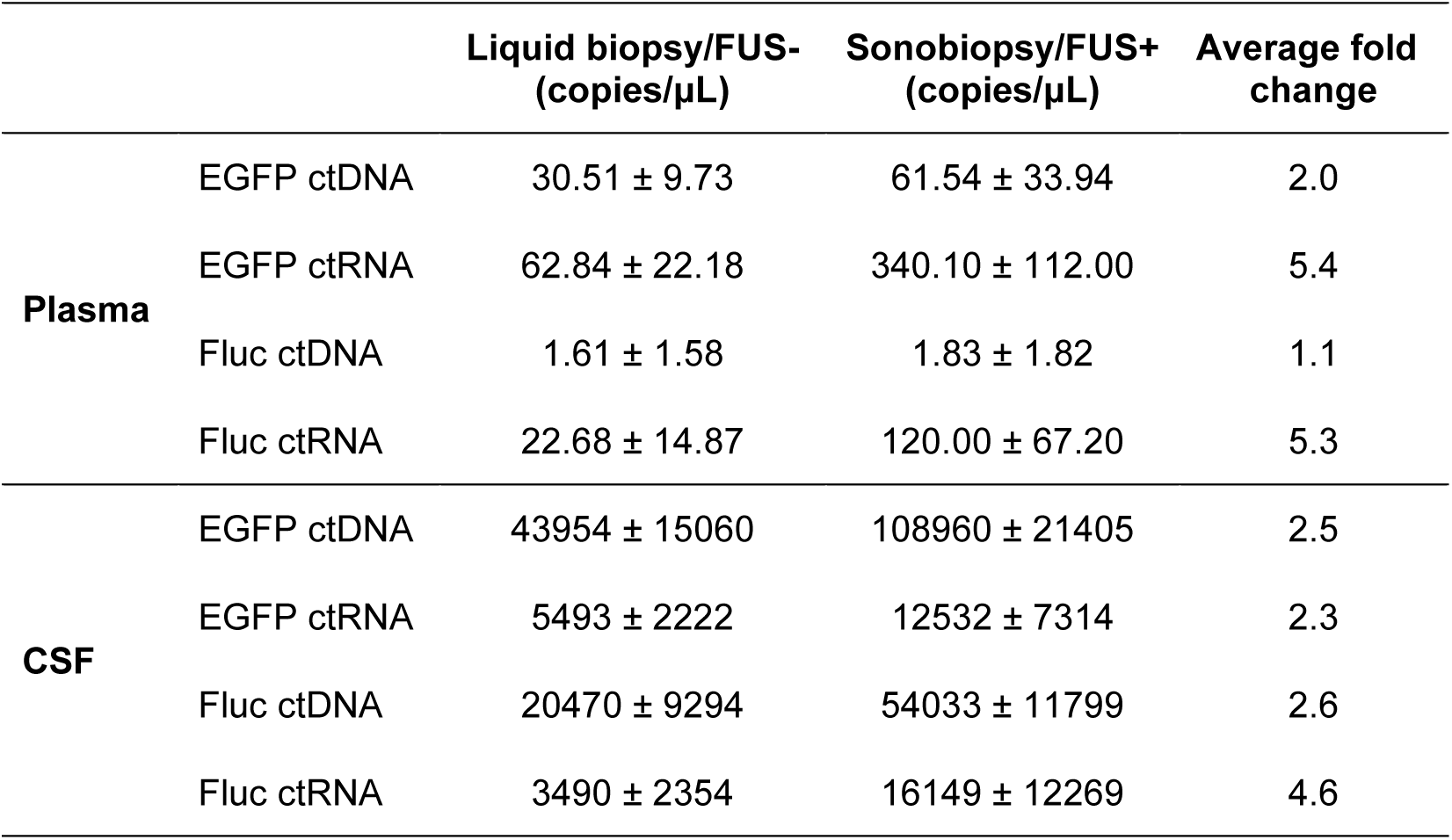
ddPCR analysis of circulating biomarkers in the plasma and CSF of a DMG mouse model.

| | | Liquid biopsy/FUS-<br>(copies/μL) | Sonobiopsy/FUS+<br>(copies/μL) | Average fold<br>change |
| --- | --- | --- | --- | --- |
| Plasma | EGFP ctDNA | 30.51 ± 9.73 | 61.54 ± 33.94 | 2.0 |
| | EGFP ctRNA | 62.84 ± 22.18 | 340.10 ± 112.00 | 5.4 |
| | Fluc ctDNA | 1.61 ± 1.58 | 1.83 ± 1.82 | 1.1 |
| | Fluc ctRNA | 22.68 ± 14.87 | 120.00 ± 67.20 | 5.3 |
| CSF | EGFP ctDNA | 43954 ± 15060 | 108960 ± 21405 | 2.5 |
| | EGFP ctRNA | 5493 ± 2222 | 12532 ± 7314 | 2.3 |
| | Fluc ctDNA | 20470 ± 9294 | 54033 ± 11799 | 2.6 |
| | Fluc ctRNA | 3490 ± 2354 | 16149 ± 12269 | 4.6 |

### FUS did not induce detectable tissue damage

As shown by the representative images of the H&E-stained brain slice, no red blood cell extravasation was observed at the FUS-targeted brain location **(Figure 6A)**. No evident microhemorrhage density change was found between the FUS-treated tumor and the non-treated tumor (*P* = .2149; unpaired *t* test) **(Figure 6B)**.

**Figure 6.**
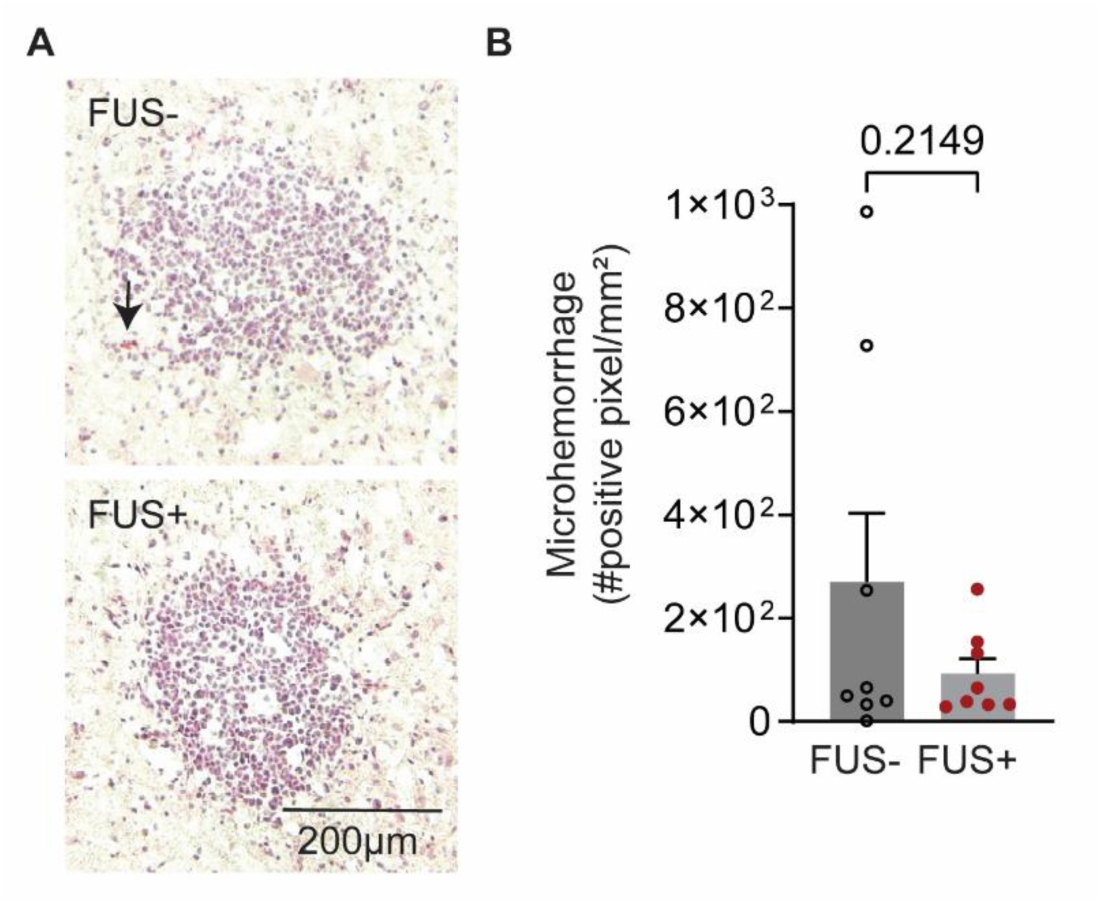
FUS did not cause significant acute tissue damage. **A.** Represent H&E staining for two subjects treated without and with FUS. The arrow indicates an example of microhemorrhage. **B.** Quantification of microhemorrhage density in the FUS- and FUS+ group. The bar graph represents the mean ± standard error of the mean (SEM). Unpaired t-test, comparing FUS+ with FUS-.

## Discussion

This study demonstrated that FUS enriched circulating biomarkers in plasma and CSF without causing brain tissue damage in a DMG mouse model. Findings from this study support the future application of sonobiopsy for noninvasive and sensitive molecular diagnosis of DMG.

ddPCR is one of the most sensitive methods for detecting tumor mutations^29^. This study developed ddPCR assays for detecting circulating tumor biomarkers for two analytes (ctDNA vs. ctRNA) in two biofluids (plasma vs. CSF) with a priori knowledge of two foreign proteins (EGFP vs. Fluc) expressed by the implanted DMG tumors. FUS opened the BBB at the spatially targeted brain location, facilitated the release of tumor-derived biomarkers from precisely defined tumor locations into the blood circulation, and enabled the timely detection of biomarkers in the blood to minimize clearance. The current study demonstrated that FUS could be integrated with ddPCR assays to assess the potential of sonobiopsy to enrich circulating biomarkers in the setting of DMG.

The effectiveness of FUS in enhancing the detection of circulating tumor biomarkers was found to depend on the source, the analytes, and the individual biomarker. FUS notably elevated the plasma levels of EGFP ctRNA by 5.4-fold and Fluc ctRNA by 5.3-fold. Additionally, FUS slightly increased the plasma level of EGFP ctDNA (but not Fluc ctDNA). These results highlight the effectiveness of FUS in enriching different circulating DMG tumor biomarkers in the plasma, partially aligning with our previous sonobiopsy studies in preclinical and clinical GBM models^7,8^. On the other hand, we found that CSF contained much higher levels of both EGFP ctDNA and Fluc ctDNA than those in plasma, consistent with previous findings on DMG liquid biopsy ^30–32^. Furthermore, our results revealed, for the first time, that FUS markedly elevated the CSF levels of EGFP ctDNA by 2.5-fold and Fluc ctDNA by 2.6-fold. These results suggest that FUS in combination with microbubbles increased BBB permeability, thereby promoting the release of tumor biomarkers from the brain into the bloodstream. Additionally, our findings indicate that FUS with microbubbles may enhance the transport of biomarkers from the interstitial fluid to the CSF. A recent study from our group demonstrated that FUS with microbubbles promotes tracer movement within the interstitial space^33^. Building upon this finding, we hypothesize that FUS with microbubbles may enrich CSF biomarkers by enhancing glymphatic transport. Nonetheless, the underlying mechanisms remain poorly understood, and further investigation is warranted.

While our study has yielded valuable insights, several limitations warrant consideration. Firstly, our investigation examined only one ultrasound parameter within this tumor model. Future investigations should encompass a broader array of FUS parameters to optimize the release of biomarkers effectively. Secondly, EGFP and Fluc were surrogate tumor biomarkers but not intrinsic DMG tumor biomarkers. Subsequent research endeavors should evaluate intrinsic DMG biomarkers, especially H3K27M mutation^34,35^. However, the H3K27M mutation was not present in our DMG cell line, preventing us from assessing the effect of FUS on this specific mutation. Future work is needed to use H3K27M-expressing DMG cells to establish the DIPG mouse model for evaluating FUS’s efficacy in enhancing the detection of H3K27M mutation and other DMG tumor biomarkers in circulation. Thirdly, the current study did not include within-subject comparisons of blood sampling before and after the FUS procedure. Collecting blood from the same subjects before and after FUS procedure within FUS+ group would enable us to use pre-FUS samples as within-subject control for evaluating the direct effect of FUS on biomarker release. Similarly, conducting two blood collections within FUS-group would allow us to evaluate any biomarker change unrelated to the FUS procedure. However, mice have a limited blood volume (∼1.5 mL). The IACUC guidelines recommend collecting no more than 10% of the total blood volume (∼150 µL) in a single week. This volume typically yields ∼75 µL of plasma, which is insufficient for generating reliable ddPCR results, as the recommended plasma input is 200 µL. Additionally, survival blood collection often results in higher rates of red blood cell lysis and increased variance between subjects compared to terminal blood collection. Therefore, in this proof-of-concept study, we employed terminal cardiac puncture to collect ∼500 µL blood samples from each mouse to ensure reliable ddPCR analysis. Lastly, future studies are needed to understand the mechanism underlying the dependency of sonobiopsy efficacy on the source, the analytes, and the individual biomarkers.

In conclusion, this study reported that FUS enriched DMG tumor biomarkers in the blood circulation and CSF. This technique offers a safe and noninvasive approach to enhance the sensitivity of liquid biopsy for DMG diagnosis. This study supported further development of this promising technique to achieve noninvasive, sensitive molecular diagnosis of DMG.

## Acknowledgments

The authors would like to thank Dr. James D. Quirk for his help with MRI.

## Authorship

J.Y. established the tumor mouse model. D.Z. and J.Y. performed mouse sonobiopsy experiments. J.Y. performed ddPCR assays for plasma and CSF. Y.Y. performed the whole brain slicing and histological staining. D.Z. and J.Y. analyzed the data. D.Z., J.Y. and H.C. wrote the manuscript with input from all other authors. H.C. supervised the research. All authors read and approved the final manuscript.

## References

1. Hargrave D, Bartels U, Bouffet E. Diffuse brainstem glioma in children critical review of clinical trials. Lancet Oncol. 2006; 7:241–248.

2. Mathew RK, Rutka JT. Diffuse intrinsic pontine glioma: Clinical features, molecular genetics, and novel targeted therapeutics. J Korean Neurosurg Soc. 2018;61(3):343–351.

3. Leach JL, Roebker J, Schafer A, et al. MR imaging features of diffuse intrinsic pontine glioma and relationship to overall survival: Report from the International DIPG Registry. Neuro Oncol. 2020;22(11):1647–1657.

4. Lu VM, Power EA, Zhang L, Daniels DJ. Liquid biopsy for diffuse intrinsic pontine glioma: An update. J Neurosurg Pediatr. 2019;24(5):593–600.

5. De Mattos-Arruda L, Mayor R, Ng CKY, et al. Cerebrospinal fluid-derived circulating tumour DNA better represents the genomic alterations of brain tumours than plasma. Nat Commun. 2015;6: 8839–8844.

6. Wei X, Meel MH, Breur M, Bugiani M, Hulleman E, Phoenix TN. Defining tumor-associated vascular heterogeneity in pediatric high-grade and diffuse midline gliomas. Acta Neuropathol Commun. 2021;9(1):142–159.

7. Meng Y, Pople CB, Suppiah S, et al. MR-guided focused ultrasound liquid biopsy enriches circulating biomarkers in patients with brain tumors. Neuro Oncol. 2021;23(10):1789–1797.

8. Yuan J, Xu L, Chien CY, et al. First-in-human prospective trial of sonobiopsy in high-grade glioma patients using neuronavigation-guided focused ultrasound. NPJ Precis Oncol. 2023;7(1):92–102.

9. Pacia CP, Yuan J, Yue Y, et al. Sonobiopsy for minimally invasive, spatiotemporally-controlled, and sensitive detection of glioblastoma-derived circulating tumor DNA. Theranostics. 2022;27(1):362–378.

10. Gorick CM, Breza VR, Nowak KM, et al. Applications of focused ultrasound-mediated blood-brain barrier opening. Adv Drug Deliv Rev. 2022;191:114583–114602

11. Pagès MDS, Rotem D, Gydush G, et al. Liquid biopsy detection of genomic alterations in pediatric brain tumors from cell-free DNA in peripheral blood, CSF, and urine. Neuro Oncol. 2022;24(8):1352–1363.

12. Meng Y, Reilly RM, Pezo RC, et al. MR-Guided Focused Ultrasound Enhances Delivery of Trastuzumab to Her2-Positive Brain Metastases. 2021;13(1):4011–4018

13. Lipsman N, Meng Y, Bethune AJ, et al. Blood–brain barrier opening in Alzheimer’s disease using MR-guided focused ultrasound. Nat Commun. 2018;9(1):2336–2343

14. Rezai AR, Ranjan M, D’haese PF, et al. Noninvasive hippocampal blood−brain barrier opening in Alzheimer’s disease with focused ultrasound. Proc. Natl. Acad. Sci. U.S.A. 2020;117(17):9180–9182

15. Abrahao A, Meng Y, Llinas M, et al. First-in-human trial of blood–brain barrier opening in amyotrophic lateral sclerosis using MR-guided focused ultrasound. Nat Commun. 2019;10(1): 4373–4381

16. Gasca-Salas C, Fernández-Rodríguez B, Pineda-Pardo JA, et al. Blood-brain barrier opening with focused ultrasound in Parkinson’s disease dementia. Nat Commun. 2021;12(1):779–785.

17. Tazhibi M, McQuillan N, Wei HJ, et al. Focused ultrasound-mediated blood–brain barrier opening is safe and feasible with moderately hypofractionated radiotherapy for brainstem diffuse midline glioma. J Transl Med. 2024;22(1):320–332

18. Hart E, Bianco J, Bruin MAC, et al. Radiosensitisation by olaparib through focused ultrasound delivery in a diffuse midline glioma model. Journal of Controlled Release. 2023;357:287–298.

19. Englander ZK, Wei HJ, Pouliopoulos AN, et al. Focused ultrasound mediated blood–brain barrier opening is safe and feasible in a murine pontine glioma model. Sci Rep. 2021;11(1): 6521–6530.

20. Haumann R, Bianco JI, Waranecki PM, et al. Imaged-guided focused ultrasound in combination with various formulations of doxorubicin for the treatment of diffuse intrinsic pontine glioma. Transl Med Commun. 2022;7(1):8–19

21. Zhang X, Ye D, Yang L, et al. Magnetic Resonance Imaging-Guided Focused Ultrasound-Based Delivery of Radiolabeled Copper Nanoclusters to Diffuse Intrinsic Pontine Glioma. ACS Appl Nano Mater. 2020;3(11):11129–11134.

22. Ishida J, Alli S, Bondoc A, et al. MRI-guided focused ultrasound enhances drug delivery in experimental diffuse intrinsic pontine glioma. Journal of Controlled Release. 2021;330:1034–1045.

23. Martinez P, Nault G, Steiner J, et al. MRI-guided focused ultrasound blood-brain barrier opening increases drug delivery and efficacy in a diffuse midline glioma mouse model. Neurooncol Adv. 2023;5(1):1–14

24. Misuraca KL, Cordero FJ, Becher OJ. Pre-clinical models of diffuse intrinsic pontine glioma. Front Oncol. 2015;5:172–178.

25. Hennika T, Hu G, Olaciregui NG, et al. Pre-clinical study of panobinostat in xenograft and genetically engineered murine diffuse intrinsic pontine glioma models. PLoS One. 2017;12(1):e0169485–e0169504

26. Choi JJ, Pernot M, Small SA, Konofagou EE. Noninvasive, transcranial and localized opening of the blood-brain barrier using focused ultrasound in mice. Ultrasound Med Biol. 2007;33(1):95–104.

27. Rizk AA, Plitman E, Senthil P, Venkatraghavan L, Chowdhury T. Effects of Anesthetic Agents on Blood Brain Barrier Integrity: A Systematic Review. Canadian Journal of Neurological Sciences. 2023;50(6):897–904.

28. Cao Y, Ni C, Li Z, et al. Isoflurane anesthesia results in reversible ultrastructure and occludin tight junction protein expression changes in hippocampal blood-brain barrier in aged rats. Neurosci Lett. 2015;587:51–56.

29. Zhang R, Han J, Daniels D, Huang H, Zhang Z. Detecting the H3F3A mutant allele found in high-grade pediatric glioma by real-time PCR. J Neurooncol. 2016;126(1):27–36.

30. Martínez-Ricarte F, Mayor R, Martínez-Sáez E, et al. Molecular diagnosis of diffuse gliomas through sequencing of cell-free circulating tumor DNA from cerebrospinal fluid. Clinical Cancer Research. 2018;24(12):2812–2819.

31. Izquierdo E, Proszek P, Pericoli G, et al. Droplet digital PCR-based detection of circulating tumor DNA from pediatric high grade and diffuse midline glioma patients. Neurooncol Adv. 2021;3(1):1–12

32. Bonner ER, Harrington R, Eze A, et al. Circulating tumor DNA sequencing provides comprehensive mutation profiling for pediatric central nervous system tumors. NPJ Precis Oncol. 2022;6(1):63–68.

33. Gong Y, Xu K, Ye D, et al. In vivo two-photon microscopy imaging of focused ultrasound-mediated glymphatic transport in the mouse brain. J Cereb Blood Flow Metab. 2025;45(7):1281–1292

34. Panditharatna E, Kilburn LB, Aboian MS, et al. Clinically relevant and minimally invasive tumor surveillance of pediatric diffuse midline gliomas using patient-derived liquid biopsy. Clinical Cancer Research. 2018;24(23):5850–5859.

35. Stallard S, Savelieff MG, Wierzbicki K, et al. CSF H3F3A K27M circulating tumor DNA copy number quantifies tumor growth and in vitro treatment response. Acta Neuropathol Commun. 2018;6(1):80–83.

